# Precise tuning of gene expression output levels in mammalian cells

**DOI:** 10.1101/352377

**Authors:** Yale S. Michaels, Mike B. Barnkob, Hector Barbosa, Toni A. Baeumler, Mary K. Thompson, Violaine Andre, Huw Colin-York, Marco Fritzsche, Uzi Gileadi, Hilary M. Sheppard, David J.H.F. Knapp, Thomas A. Milne, Vincenzo Cerundolo, Tudor A. Fulga

**Author notes:** Correspondence should be addressed to T.A.F.

## Abstract

Precise, analogue regulation of gene expression is critical for development, homeostasis and regeneration in mammals. In contrast, widely employed experimental and therapeutic approaches such as knock-in/out strategies are more suitable for binary control of gene activity, while RNA interference (RNAi) can lead to pervasive off-target effects and unpredictable levels of repression. Here we report on a method for the precise control of gene expression levels in mammalian cells based on engineered, synthetic microRNA response elements (MREs). To develop this system, we established a high-throughput sequencing approach for measuring the efficacy of thousands of miR-17 MRE variants. This allowed us to create a library of <u>mi</u>croRNA <u>s</u>ilencing-mediated <u>f</u>ine-<u>t</u>uners (miSFITs) of varying strength that can be employed to control the expression of user specified genes. To demonstrate the value of this technology, we used a panel of miSFITs to tune the expression of a peptide antigen in a mouse melanoma model. This analysis revealed that antigen expression level is a key determinant of the anti-tumour immune response *in vitro* and *in vivo*. miSFITs are a powerful tool for modulating gene expression output levels with applications in research and cellular engineering.

## INTRODUCTION

Subtle changes in gene expression can have important biological consequences^1–3^. To explore the impact of partial changes in gene expression, fine-tuning systems based on libraries of promoters or ribosome binding sites of varying strengths have previously been constructed in bacteria^4–6^ and yeast^4, 7^. Here, we set out to develop a tool that would enable precise, stepwise modulation of gene expression levels in mammalian cells. To create a generalizable gene-tuning technology and overcome common limitations of existing genetic manipulation methods we aimed to design a system which: *i)* is free from antibiotic triggers, such as doxycycline or rapamycin^8, 9^; *ii)* does not rely on introducing exogenous siRNAs as these can induce broad off-target effects^10^; and *iii)* is not dependent on artificial promotors or upstream open reading frames that may not be portable to all proteins or cell types due to the highly context-dependent nature of gene regulation in mammals^11^.

To satisfy these design criteria, we sought to harness the exquisite ability of microRNAs (miRNAs) to fine-tune gene expression in mammalian cells. miRNAs are short non-coding RNAs capable of post-transcriptionally controlling gene expression levels by recruiting the RNA induced silencing complex (RISC) to cellular RNAs bearing cognate miRNA response elements (MREs). Importantly, the magnitude of repression depends on the extent of complementarity between a miRNA and its target MRE^12^. We reasoned that by engineering synthetic MREs with varying complementarity to an endogenous miRNA we could precisely modulate expression of user-specified genes without the necessity of supplying any exogenous molecules.

Previous high-throughput screening approaches have enabled in depth analysis of miRNA expression profiles^13^ and the evaluation of contextual features important for miRNA-mediated regulation^14^. Additional studies have described broad functional domains within MREs, such as the “seed” (nt 2-8) and the “supplementary region” (nt 13-16)^12^. However, it remains unclear how each individual nucleotide or pair of nucleotides within a MRE contributes to the degree of gene silencing imparted by a given miRNA in living cells. Furthermore, although miRNAs have been reported to promote translational repression ^15–17^ in addition to degradation of mRNA targets, it is unknown how the complementarity between a miRNA and its cognate MRE influences each of these mechanisms.

To further our understanding of miRNA-MRE interactions and to enable the forward design of a gene-tuning technology, we developed a high-throughput approach to dissect the functional landscape of a MRE at single-base resolution. We identified nucleotides that are critical for repression and determined that miRNA-MRE complementarity dictates transcript degradation as well as translational repression. We then used this information to create a panel of <u>mi</u>RNA <u>s</u>ilencing-mediated <u>fi</u>ne-<u>t</u>uners (miSFITs) that allowed us to precisely modulate the expression levels of PD-1, a T-cell inhibitory receptor and an important target for cancer immunotherapy. Finally, we employed the miSFIT approach to fine-tune a tumour-associated antigen in a melanoma model. This allowed us to identify antigen expression level as an important determinant of the anti-tumour immune response *in vitro* and *in vivo*.

## RESULTS

### Dissecting the regulatory landscape of a miRNA response element

To develop a fine-tuning system suitable for use in mammalian cells, we sought to redirect endogenous miRNAs to user-defined target mRNAs, thus harnessing the repressive potential of this post-transcriptional regulatory layer. As a proof of concept, we focused on miR-17 which is a well characterized miRNA expressed in numerous human and murine cell types^18^. By evaluating the regulatory capacity of a library of MREs with varying complementarity to miR-17 we reasoned that we could dissect the targeting landscape of this miRNA. The resulting dataset could be used to select MREs of desired strength, providing a generalizable approach for fine-tuning gene expression.

We designed a 23nt degenerate oligonucleotide pool with 91% complementarity to miR-17 and 3% of each alternative nucleotide at every position (Fig. 1a, Supplementary Fig. 1). This oligo pool was cloned downstream of a fluorescent reporter (ECFP) in a mammalian expression plasmid and the ensuing MRE variant library was transfected into HEK-293T cells that endogenously express miR-17. We also co-transfected a control reporter bearing an MRE complementary to *C. elegans* Cel-miR-67, which is not expressed in human cells^19^. After allowing endogenous miR-17 to act on the transcripts templated by the variant library, we harvested mRNA and plasmid DNA (pDNA) and subjected them to targeted deep sequencing (Fig. 1b, Supplementary Fig. 1). To estimate the strength of the MRE variants present in our library, we divided their frequency in the mRNA pool by their frequency in the pDNA pool (Supplementary Fig. 1).

**Figure 1.**
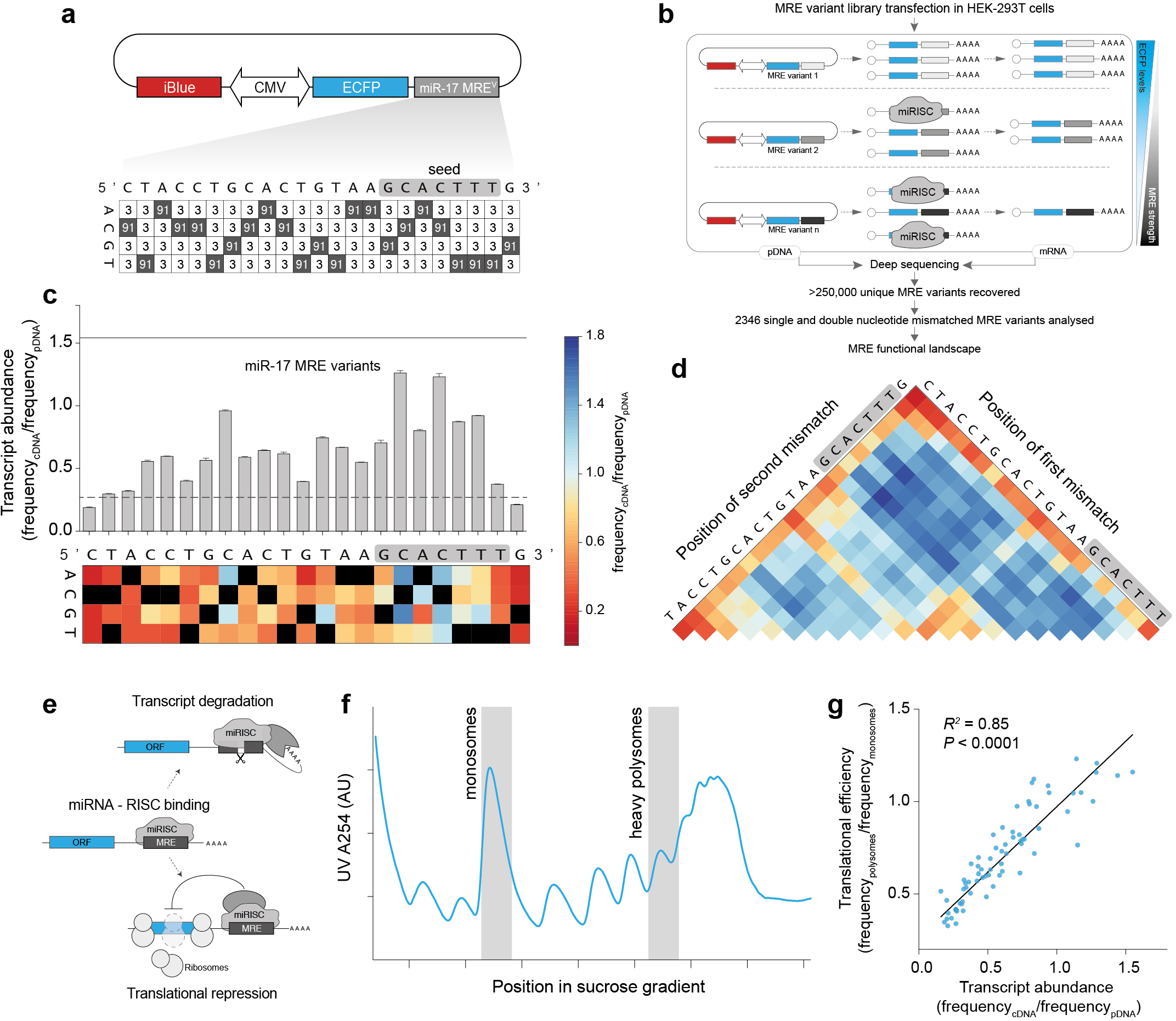
Analysis of MRE regulatory landscape at single-nucleotide resolution. **(a)** MRE reporter library diagram. Values indicate the proportion of nucleotides at each position in the MRE (shaded squares = nucleotides complementary to miR-17). **(b)** MRE regulatory landscape analysis pipeline. **(c)** Impact of MRE variants on transcript abundance. Bar graph shows relative contribution of each nucleotide to MRE function, as determined by high-throughput sequencing (n = 3 biological replicates, mean +/− s.d.; dashed line = expression of a perfectly complementary MRE, solid line = expression of a non-targeted MRE-Cel-miR-67). Heat-map displays the effect of each possible mismatch by position and reflects the mean of three replicates (complementary bases are displayed in black). **(d)** The impact of di-nucleotide substitutions on reporter expression (mean of 3 biological replicates; colour scale is the same as in (c); grey box = seed region). **(e)** Schematic representation of the two major pathways underlying miRNA-mediated repression. **(f)** Polysome profiles generated by sucrose gradient fractionation. Blue trace denotes the spatial distribution of RNA across the gradient as monitored by UV absorbance. Analysed fractions (monosomes and heavy polysomes) are shaded in grey (representative of two biological replicates). **(g)** Correlation between translational efficiency and transcript stability for all single nucleotide miR-17 MRE variants. The slope of a linear regression model (black diagonal line) significantly differs from 0.

As expected, MREs with higher complementarity to miR-17 were silenced more effectively (Supplementary Fig 1). Even single nucleotide mismatches diminished silencing by 2.30-fold on average (+/− 0.03, 95% CI) compared to a perfectly matched target (Supplementary Fig. 1). We then focused our analysis on all single nucleotide variants and asked how each position within the MRE contributes to miRNA-mediated repression (Fig. 1c). As anticipated, certain seed mismatches strongly abrogated silencing, confirming the important role of this region in target selection (Fig. 1c). Intriguingly however, non-seed nucleotides also significantly impacted the degree of repression, with one position even having a greater impact on silencing than most seed nucleotides (Fig. 1c). Mutations introducing G:U wobble pairs were always less deleterious to silencing than non-pairing bases, highlighting the importance of thermodynamic stability for miRNA-mediated repression (Fig. 1c). Analysis of double-nucleotide variants revealed that pairs of mismatches within the seed or combinations of seed mismatches with mismatches in positions 14 to 20 strongly impaired miRNA activity (Fig. 1d). When we subjected a second miRNA (miR-21) to the same high-resolution analysis, the relative importance of each position in the MRE correlated only weakly with miR-17 (*R*^2^ = 0.22, *P* = 0.03) (Supplementary Fig. 2). Notably however, despite this weak correlation, mismatches at certain non-seed positions were also able to strongly abrogate silencing by miR-21. Together, these data demonstrate the utility of our high-throughput assay for studying the functional landscape of MREs at single nucleotide resolution and reveal miRNA specific targeting preferences that may not be predicted by generic algorithms^20^. These results also suggest that choosing other input miRNAs for tuning gene-expression may require additional empirical analysis.

This sequencing-based assay allows us to assess the effect of miRNA / MRE mismatches on mRNA stability. In addition to promoting transcript degradation, miRNAs have also been proposed to repress translation^21^. However, the extent to which these two processes are correlated across different MREs remains unclear^15–17^ (Fig. 1e). To determine the degree to which the MREs in our library mediate translational repression we used polysome profiling to isolate monosome-bound and heavy polysome-bound mRNAs^22^ (Fig. 1f). We then sequenced cDNA libraries from these fractions and used the ratio of reads in the heavy polysome-bound fraction to reads in the monosome-bound fraction as a measure of translational efficiency for each MRE variant (Fig. 1f, g). This analysis revealed a strong correlation between transcript degradation and translational repression (the inverse of translational efficiency) for single-nucleotide variants in the library (Fig. 1g) (*R*^2^ = 0.85, *P* < 0.0001, linear regression). This finding suggests that miRNA-target base pairing is a critical determinant of the magnitude of both transcript degradation and translational repression. These data also indicate that our mRNA/pDNA sequencing approach is a good predictor of overall MRE strength.

To further validate the accuracy of our high-throughput MRE screen, we randomly selected 15 single and double nucleotide MRE variants from our library and subjected them to RT-qPCR and flow-cytometry analysis (Supplementary Fig. 3a). Both RT-qPCR (*R*^2^ = 0.92, linear regression, Supplementary Fig. 3c) and flow-cytometry (*R*^2^ = 0.95, linear regression, Supplementary Fig. 3b, d) strongly corroborated the high-throughput sequencing analysis, supporting the validity of our screen and confirming the correlation between the strength of transcript degradation and translational repression in this particular context (Supplementary Fig. 3e).

### Fine-tuning PD-1 expression in Jurkat T-cells

Next, we sought to demonstrate that our MRE variant library can be used to precisely modulate expression of a gene of interest. By ranking all miR-17 MRE variants containing single-nucleotide mismatches, we created a dictionary of <u>mi</u>croRNA <u>s</u>ilencing-mediated <u>fi</u>ne-<u>t</u>uners (miSFITs) that relates MRE sequence identity to ECFP gene expression output in HEK-293T cells (Fig. 2a). Sorting all miSIFTs according to their predicted strength revealed that the system has the capacity to achieve precise, stepwise control of gene expression levels (Fig. 2a, Supplementary Fig. 4a). More specifically, the difference in expression between adjacent single-nucleotide miSFIT variants is 1.3% of maximal expression on average (0.87% to 1.77%, 95% CI, Supplementary Fig. 4a).

**Figure 2.**
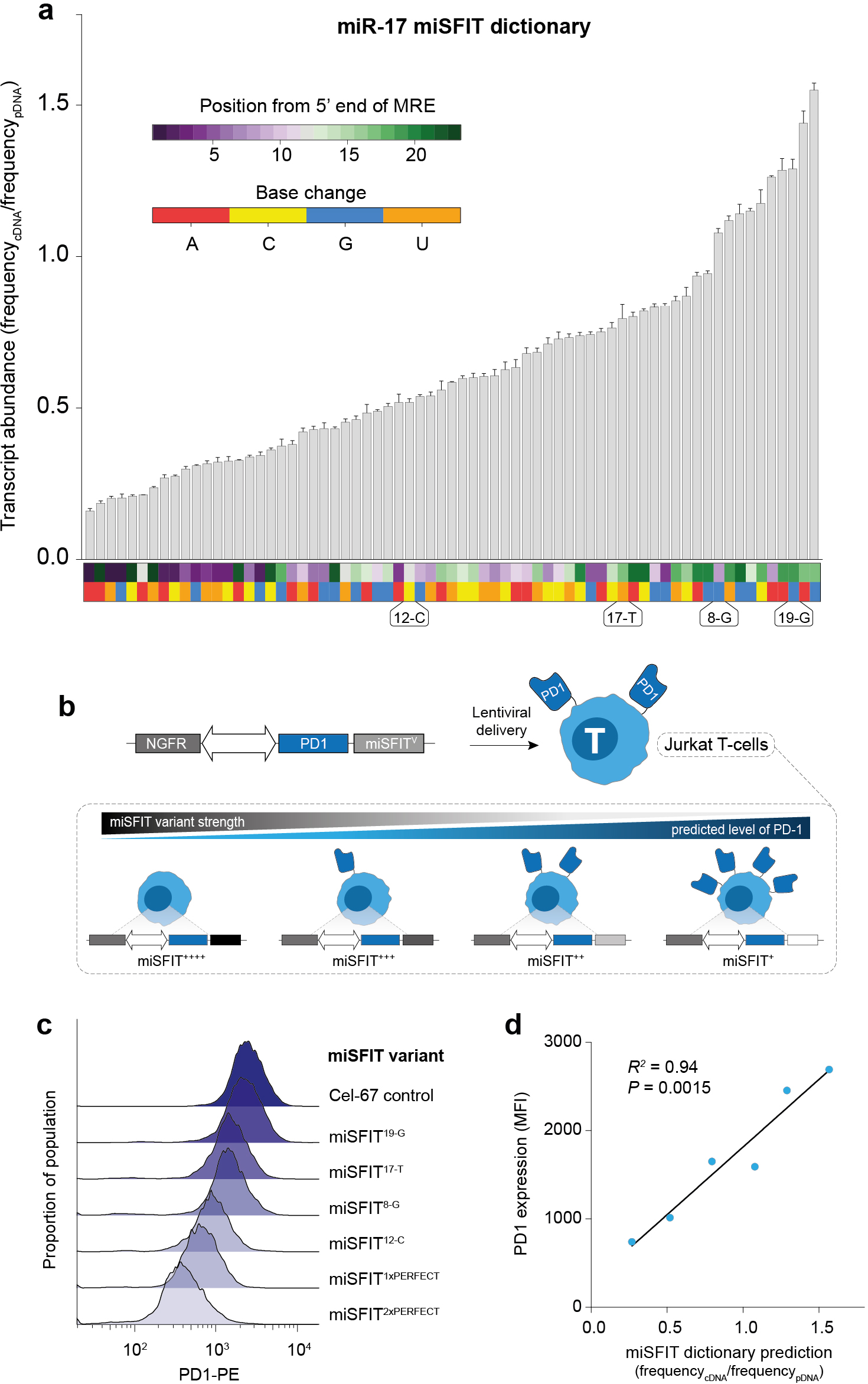
Synthetic miSFIT variants enable fine-tuning of gene expression in mammalian cells. **(a)** Impact on transcript abundance of all single-nucleotide miR-17 miSFIT variants ranked by expression output. Coloured rectangles beneath each bar indicate the position (top) and base change (bottom) of the synthetic MRE variant (n = 3 biological replicates, mean +/− s.d.; annotated positions = miSFIT variants used in (b-d)). **(b)** Schematic of miSFIT tuning strategy. PD-1 expression is controlled by various miSFIT variants while NGFR serves as an un-silenced internal control. **(c)** Flow-cytometry histograms of PD-1 expression on Jurkat T-cell lines transduced with one of six different miSFIT variants (x-axis = log_10_ transformed PD1-PE fluorescence). **(d)** Correlation between predicted expression in the miSFIT dictionary and observed PD1 expression on Jurkat T-cell lines (linear regression).

We then asked if a selection of miSFIT variants from this dictionary, could be used in a different human cell type to tune expression of a protein with an important biological function. Programmed cell death 1 (PD-1) is a co-inhibitory receptor expressed on effector T cells and an important target for anti-cancer immunotherapy^23^. We selected four miR-17 miSFITs from the ECFP dictionary (Fig 2a), a perfectly complementary MRE (1x perfect site), tandem perfectly complementary MREs (2x perfect sites), and the control miR-Cel-67 MRE. We appended each variant downstream of *PD-1* in a bi-cistronic lentiviral vector that also encodes a control reporter gene (truncated nerve growth factor receptor, NGFR) that is not under MRE control^13^ (Fig. 2b). We then transduced Jurkat T-cells, that express very low levels of PD-1 at baseline, with each of these constructs at low MOI. After sorting pools of NGFR^+^ cells we assayed PD-1 expression by flow cytometry. The selected miSFITs elicited discrete, stepwise control over PD-1 levels (Fig. 2c, Supplementary Fig. 4) in a manner that was predicted by the ECFP MRE dictionary (Fig. 2d, *R*^2^ = 0.94, linear regression).

### Modulating tumour associated antigen expression and T-cell response

To further illustrate the utility of miSFITs as an effective tool for modulating gene-expression, we next sought to apply this technology towards a biological question that has previously been confounded by technical limitations. More specifically, we set out to explore how peptide-antigen expression levels influence the strength of the anti-tumour immune response in a murine melanoma model. Cancer immunotherapy is a promising class of treatments that aim to enhance anti-tumour cytotoxicity by the adaptive immune system^24^. Sub-types of immunotherapy, including checkpoint blockade and adoptive cell transplant, rely on T-cell receptor (TCR) mediated recognition of peptide antigens presented by MHC-I molecules on the surface of tumour cells^24^. Although *in silico* algorithms can accurately predict which peptide antigens are likely to elicit an immune response^25^, understanding how peptide-antigen expression levels influence the strength of the anti-tumour immune response *in vivo* remains elusive. A quantitative analysis of this relationship could provide an important benchmark for predicting which tumours might respond to anti-cancer immunotherapy.

Previous efforts to titrate peptide-MHC concentrations have relied on coating culture vessels with recombinant peptide-MHC multimers^26^ or by briefly adding varying concentrations of peptide to cellular growth media (a process known as peptide pulsing)^27^. Although valuable, these methods cannot accurately re-capitulate the endogenous pathway of antigen expression, proteolytic processing and subsequent surface presentation. Furthermore, because peptide pulsing is inherently transient, this method precludes tracking the survival of antigen-expressing cells *in vivo*. To understand how antigen-expression influences the anti-tumour immune response and the relative fitness of cancer cells *in vitro* and *in vivo*, we used miSFITs to finely tune expression of ovalbumin (OVA), a model immunogenic protein, in a stable and physiologically accurate fashion.

To this end, we created a panel of seven bi-cistronic OVA expression vectors, each encoding a distinct miSFIT variant in the 3’UTR of ovalbumin (Fig. 3a). We also coupled EGFP downstream of ovalbumin *via* a self-cleaving T2A peptide, enabling us to monitor expression levels by flow-cytometry (Fig. 3a). In each vector, NGFR was included as an unsilenced internal control reporter. We transiently expressed these constructs in B16-F10 melanoma cells to evaluate gene expression output (Supplementary Fig. 5a). This analysis revealed discrete, stepwise tuning of target levels, although the exact ranking of miSFIT variant strength differed from what we observed when tuning PD-1 in human Jurkat T-cells (Supplementary Fig. 5b). To generate stable cell lines expressing varying levels of ovalbumin we then transduced B16-F10 cells with a subset of five OVA-miSFIT constructs at low MOI (<2% transduction efficiency). The semi-random nature of lentiviral integration^28^ results in heterogeneity of gene expression between individual cells. To mitigate this effect, we sorted and expanded pools of 150,000 cells on the basis of NGFR expression. After confirming that we successfully tuned ovalbumin expression in the resulting five cell lines (Fig. 3b), we asked how antigen expression levels influence CD8^+^ T-cell activation.

**Figure 3.**
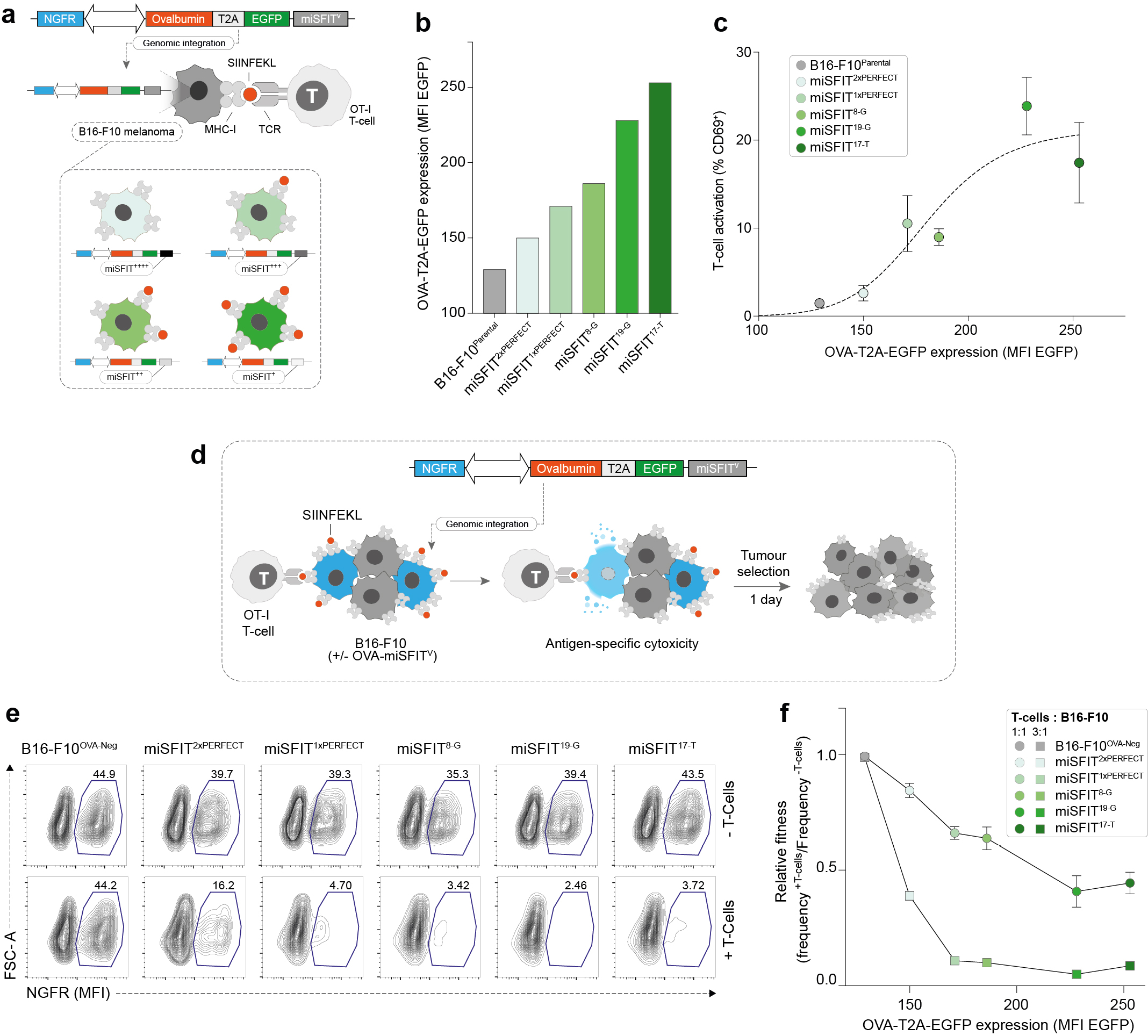
Fine-tuning antigen expression levels and T-cell activity. **(a)** Strategy for tuning Ovalbumin (OVA) expression in B16-F10 melanoma cells using lentivirally integrated miSFITs. Red circles represent SIINFEKL, a peptide antigen derived from ovalbumin. OT-I T-cells express a TCR specific for SIINFEKL presented on MHC-I. **(b)** Flow cytometry analysis of OVA-T2A-EGFP expression in B16-F10 cell lines transduced with six different miSFIT variant (miSFIT^V^) lentiviruses (see Supplementary Fig. 5 for the gating strategy and distribution of fluorescence intensity) **(c)** CD8^+^, OT-I T-cell activation by OVA-miSFIT B16-F10 cell lines. CD69 expression was quantified by flow-cytometry (n = 5 biological replicates, mean +/− s.d.). **(d)** Schematic representation of mixed-culture experimental design. OVA-negative (NGFR^−^) are mixed with OVA-miSFIT (NGFR^+^) B16-F10 cells and are challenged overnight with OT-I T-cells. **(e)** Representative flow cytometry plots of mixed culture experiments. The percentage of NGFR^+^ (OVA-miSFIT) cells (blue polygon gate) surviving after overnight selection in the presence or absence of CD8^+^, OT-I T-cells(at a ratio of 3:1 T-cells to B16-F10 cells) is indicated for each condition. **(f)** Relative fitness of B16-F10 cell lines as a function of OVA expression. Relative fitness was calculated by dividing the frequency of NGFR^+^ cells with T-cells by the frequency of NGFR^+^ cells without T-cells (n = 3 biological replicates, mean +/− s.d.).

The OT-I T-cell receptor (OT-I) is specific for SIINFEKL, a short peptide antigen derived from ovalbumin, presented by MHC-I^29^. We co-cultured each of the five B16-F10 lines expressing differential ovalbumin levels and the OVA-negative parent line with CD8^+^ OT-I T-cells and assayed activation by measuring CD69 expression (Fig. 3c). Indeed, increasing OVA expression resulted in a concomitant increase in the proportion of activated T-cells, presumably due to the greater probability of each T-cell encountering and responding to a SIINFEKL-MHC-I complex (Fig. 3c).

Under selective pressure by the adaptive immune system, tumours have been shown to acquire mutations that prevent effective T-cell surveillance in a process known as immunoediting. This is generally achieved through loss of function mutations in MHC genes, up-regulation of immunosuppressive molecules or by elimination of clones expressing neo-antigens^30^. In addition to these reported phenomena, we hypothesized that tumour cells might also be selected on the basis of antigen expression levels. To address this question, we first mixed the five OVA-miSFIT B16-F10 cell lines at a 1:1 ratio with OVA-negative B16-F10 cells (Fig. 3d). We then allowed these mixed cultures to grow overnight in the presence or absence of OT-I T-cells. Because all OVA-miSFIT lines express NGFR whilst the OVA-negative parent line does not, we quantified the relative abundance of OVA^+^ (NGFR^+^) and OVA-negative (NGFR^−^) cells following the T-cell challenge (Fig. 3e, Supplementary Fig. 5d, e). Tuning antigen expression using miSFITs modulated the strength of T-cell mediated selection in a dose-responsive manner at two T-cell: tumour cell ratios (Fig. 3e, f). Notably, even low antigen expression was sufficient to elicit a strong reduction in relative fitness at a high T-cell: tumour cell ratio (Fig. 3f).

### Antigen expression level is a key determinant of tumour growth and survival

Next, we asked if the effect of antigen expression on melanoma survival that we observed *in vitro* correlates with tumour growth rates *in vivo*. First, we injected a subset of our engineered OVA-miSFIT-B16-F10 cell lines into syngeneic recipient mice (Fig. 4a). After allowing intradermal tumours to establish for seven days, we adoptively transferred CD-8^+^ OT-I T-cells and monitored tumour growth for an additional 22 days (Fig. 4a). Antigen expression levels significantly impacted tumour growth *in vivo* in a manner that faithfully mirrored *in vitro* T-cell activation and killing (Fig. 4b) (*P* = 0.02, Kruskal-Wallis test, comparison of tumour volumes at day 19). We continued to monitor mice for 46 days and observed that antigen expression markedly influenced survival (*P* = 0.0038, Logrank test for trend, Fig. 4c). Mice bearing tumours with no, or low antigen expression all met our endpoint criteria by day 27. In contrast, medium or high OVA expressing tumours displayed a substantial increase in survival. One third of the mice bearing high-antigen B16-F10-OVA cells (2/6) survived for 46 days with tumours that were nearly undetectable by the experiment’s endpoint (Fig. 4c). Together, these findings illustrate the importance of tumour-associated antigen expression levels in determining the strength of the immune-response.

**Figure 4.**
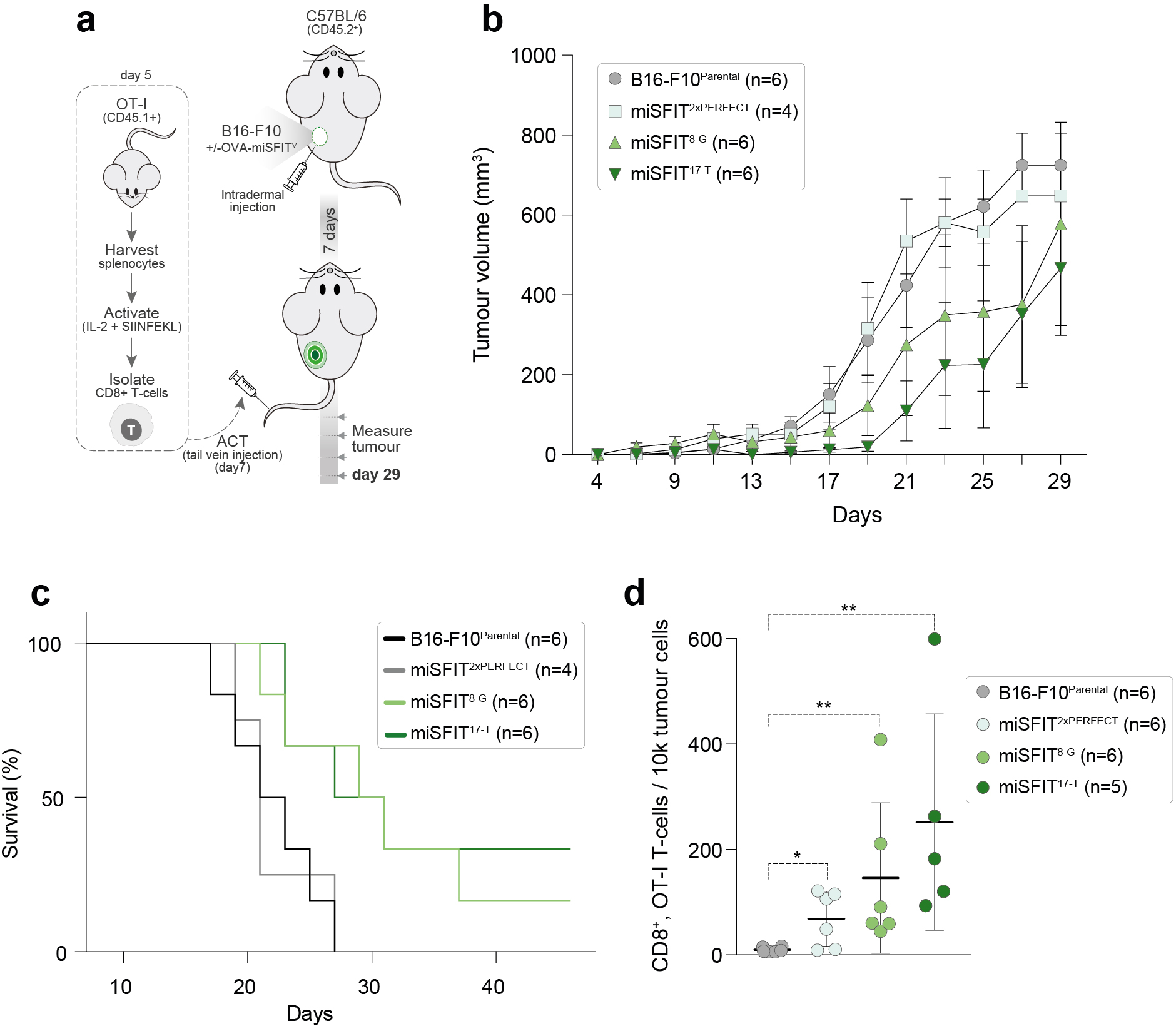
Antigen expression levels determine the anti-tumour immune response *in vivo*. **(a)** Experimental design for *in vivo* OVA-miSFIT B16-F10 tumour growth experiments. **(b)** Analysis of tumour volume over time for four B16-F10 cell lines (3 OVA-miSFIT variant lines and one B16-F10 parental control line) challenged with OT-I CD8^+^ T-cells (x-axis = number of days from tumour cells injection; mean +/− s.e.m.) **(c)** Survival curves for mice injected with B16-F10 lines following the same experimental setup as in (a). **(d)** Frequency of CD8^+^, OT-I TILs per tumour (mean +/− s.d., *Mann-Whitney U* test, \**P* < 0.05, \*\**P* < 0.01). For experimental setup see Supplementary Fig. 6a.

To understand why higher antigen expressing tumours were more effectively controlled, we harvested and analysed tumour infiltrating lymphocytes (TILs) at eight days after adoptive T-cell injections (Supplementary Fig. 6). Changes in antigen expression levels differentially affected expression of CD69, CD25, PD-1 and CTLA-4 on OT-I T-cells (Supplementary Fig. 6). Importantly, increasing levels of OVA lead to a dose-responsive increase in the frequency of TILs *in vivo* (*P* = 0.003, Kruskal-Wallis test, Fig. 4d). These findings highlight the value of miSFIT technology in studying intracellular interactions and cellular fitness, and demonstrate the role of tumour-associated antigen levels in controlling T-cell infiltration, tumour growth and survival.

## DISCUSSION

Here we have developed a powerful tool for tuning gene expression output in mammalian cells and used it to uncover a critical role of cancer antigen expression in modulating the immune response. It should be noted that the ovalbumin-derived model antigen SIINFEKL is recognized by the OT-I TCR with very high affinity. However, patient derived tumour-associated antigens have varying affinity and avidity for their cognate TCRs. Applying the miSFIT technology to bona-fide tumour antigens will enable scientists to understand how antigen immunogenicity^31^ and expression levels interact to influence the immune response. In turn, such studies could allow clinicians to better predict how tumours will respond to immunotherapy.

Although miSFITs enabled precise control of OVA expression, some of our MREs (including the Cel-miR-67 control MRE) elicited stronger or weaker silencing than expected based on the MRE dictionary (compare Fig. 2c, 3b). This context-dependent effect may be the result of differences in miR-17 family member expression or the distinct repertoire of endogenous RNA binding proteins between human (HEK 293T, Jurkat T-cells) and murine (B16-F10) cells. While variable miRNA expression profiles may necessitate cell-type specific validation of each miSFIT’s strength, this feature also presents an opportunity for context-specific gene tuning. Designing miSFIT variants with complementarity to cell-type/cell-state specific miRNAs will enable gene-tuning that is similarly confined to user-specified cell populations^32^ or that can be triggered by physiological stimuli.

The miSFIT approach displays a number of advantages over existing methods for manipulating gene-expression levels. Unlike titratable promotors, miSFITs do not require chemical inducers that have confounding effects on cellular metabolism and are difficult to dose *in vivo*^8^. Controlling viral multiplicity of infection might allow coarse control over gene expression. However, as reported in this study, even at single copy integration, strong viral promotors instigate over-expression above physiologically relevant levels. Furthermore, miSFITs’ short length makes them amenable to integration into genomic loci using CRISPR/Cas9 editing with single stranded oligonucleotide homology donors (ssODNs). This offers an opportunity to repress the expression of endogenous genes in a precise fashion. Unlike siRNAs, miSFITs co-opt endogenous miRNAs to regulate gene expression. Since the sequence space of endogenous miRNAs is several orders of magnitude smaller than that of the transcribed genome, miSFITs are not confounded by the off-target specificity issues associated with introducing exogenous siRNAs. Finally, unlike existing modalities, miSFITs have the potential to tune both over-expression of transgenes and expression of endogenous genes in a precise, stepwise fashion.

In addition to their value as a research tool, we propose that miSFITs will have future therapeutic applications. Gene-expression levels can influence the efficacy of anti-cancer immunotherapy. On tumour-reactive T-cells, high PD-1 expression suppresses effector function^23^. Therapies that block PD-1 signalling can improve the anti-tumour immune response^23^ but also instigate adverse autoimmunity events in a large proportion of patients^33^. Using miSFITs to fine-tune endogenous PD-1 levels in patient-derived effector T-cells might achieve an optimum balance between exhaustion and autoimmunity, enabling safer and more effective adoptive cell therapy. Similarly, miSFITs could be applied to other co-inhibitory receptors like CTLA-4 or to therapeutic transgenes such as Chimeric Antigen Receptors (CARs). Because of their precision and versatility, miSFITs hold promise for tuning expression of a wide range of genes with applications in basic research and therapeutic cellular engineering, complementing existing tools for binary control of gene activity.

## ONLINE METHODS

### MRE variant library construction

The sequences of all oligonucleotides used in this study are listed in Supplementary Table 1. We purchased hand-mixed, partially degenerate oligonucleotides from Integrated DNA Technologies (IDT) comprising a constant flanking region and a variable region with partial complementarity to either hsa-miR-17 or hsa-miR-21. In this study, the term synthetic MRE refers to a sequence of equal length, and largely complementary to a given miRNA. To achieve maximum coverage of single and double nucleotide variants, 91% of the complementary base and 3% of each non-complementary base were incorporate at each position in the MRE libraries. Each degenerate oligo was PCR amplified in triplicate using Phusion High-Fidelity PCR Master Mix with GC Buffer (NEB), using primers miR17_Lib_Gen_F and miR17_Lib_Gen_R, which append BsmBI recognition sites on both sides of the MRE. The resulting PCR products were pooled and purified using the MinElute PCR Purification Kit (Qiagen).

We performed a large-scale restriction cloning reaction to ligate the degenerate MRE PCR product into a reporter plasmid. The reporter plasmid comprises a bi-directional CMV promotor driving expression of iBlue fused to a degradation signal derived from Ornithine Decarboxylase and ECFP fused to the same degradation signal. We linearized 10.5 μg of the reporter plasmid downstream of ECFP by digesting with BsmBI.

The degenerate MRE PCR product (300ng) was cut with BsmBI and ligated to the linearized, dephosphorylated (Antarctic Phosphatase, NEB) and gel purified (QIAquick Gel Extraction Kit, Qiagen) reporter plasmid using T4 DNA Ligase (NEB) at 16°C overnight. We purified the ligations using the QIAquick PCR Purification Kit (Qiagen) and transformed approximately 3.6 μg of the purified product into 10-beta Electrocompetent E.coli (NEB) following the manufacturer’s instructions. Transformants were plated overnight at 32°C on 24.5 cm^2^ ampicillin-treated LB agar plates. We recovered the resulting plasmid library using the QIAfilter Plasmid Midi Kit (Qiagen).

### HEK-293T cell culture and MRE variant library transfection

HEK-293T cells were grown in Dulbecco’s modified Eagle’s medium (DMEM, Gibco) supplemented with 15% FBS (GIBCO) and 1% Penicillin-Streptomycin (P/S, 10,000 U/mL, Gibco). We screened cells for mycoplasma at the outset of the project. Cells were seeded in 12 well plates, 24 hours prior to transfection, allowing them to reach 80-90% confluency on transfection day. On the day of transfection, we replaced complete growth media with DMEM, 2% FBS (no P/S). We prepared three independent transfection mixtures, each containing 4 μg of the degenerate MRE reporter library, 4 ng of miR-Cel-67 MRE control plasmid and 12 μl Polyethylenimine (PEI, 1 mg/ml, Sigma-Aldrich) in 400 μl Opti-MEM (Gibco). Each mixture was applied dropwise to 4 wells of a 12 well plate and incubated for 24 hours.

### Polysome profiling

To generate enough cell lysate for polysome profiling, we seeded HEK-293T cells in two independent 15 cm^2^ culture dishes, allowing them to reach 70-80% confluency by the day of transfection. For each dish, we combined 25 μl each Lipofectamine 3000 and P3000 Reagent (Thermo Fisher) with 12.5 μg of the degenerate MRE reporter library and 100 ng of miR-Cel-67 MRE control plasmid, transfected according to the manufacturer’s instructions and incubated for 24 hours. To arrest translation, cycloheximide (CHX, Merck) was added to the culture dishes at 100 μg/mL for 10 minutes at 37°C. Next, dishes were placed on ice and washed with cold PBS (Life Technologies) supplemented with CHX (100μg/mL). We scraped the dishes in PBS + CHX (100μg/mL), centrifuged the harvested cells at 1,000xg for 3 minutes at 4°C and discarded the supernatant. Cell pellets were re-suspended in 200 μl of hypotonic lysis buffer (10mM HEPES pH 7.8, 1.5mM MgCl_2_, 10mM KCl, 0.5mM DTT, 1% Triton X-100 and 100mg/mL CHX) and incubated for 5 minutes on ice. Next, we lysed the cells with 10 strokes through a 26 gauge needle and pelleted the nuclei by centrifuging at 1,500xg for 5 min at 4 °C. The supernatant was flash frozen in liquid nitrogen and stored at −80 °C.

10-50% (W/V) sucrose gradients were generated using a Gradient Master (Biocomp Instruments) from 10% and 50% sucrose solutions in gradient buffer (100mM KCl, 5mM MgCl_2_, 20mM HEPES-KOH pH7.5, 1mM DTT, 100μg/mL CHX). We thawed the cell lysates and layered them on top of the chilled sucrose gradients before centrifuging at 4°C for 2 hours at 36,000 RPM in a SW-41 rotor. Gradients were fractionated from the top using a Gradient Fractionator (Biocomp Instruments). To recover RNA from the resulting fractions we added 2.25 volumes of 8M Guanidine HCl (Sigma-Aldrich) and vortexed the samples. Next, 3.25 volumes of isopropanol were added and samples were incubated overnight at −20°C. Reactions were centrifuged at > 12,000 rpm for 30 minutes at 4°C and the supernatant was aspirated. RNA pellets were re-suspended in a mixture of 90 μl nuclease free H_2_O (Invitrogen), 10 μl 3M Sodium Acetate (Invitrogen) and 1 μl 5mg/ml glycogen (Ambion) and precipitated with 250 μl of cold 100% ethanol (VWR). After 30 minutes incubation on ice, samples were centrifuged at >12,000 rpm for 30 minutes at 4°C. Next, pellets were washed with 500μl of 70% ethanol and re-suspended in nuclease free H_2_O.

### pDNA and cDNA library preparation and high-throughput sequencing

We used the All Prep DNA/RNA Mini kit (Qiagen) to simultaneously extract plasmid DNA (pDNA) and mRNA from HEK-293T cells transfected with the degenerate MRE reporter library. After performing a genomic DNA wipe-out, cDNA was generated from mRNA and polysome-associated RNA using the QuantiTect Reverse Transcription kit (Qiagen) following the manufacturer’s instructions. To create amplicon libraries for high-throughput sequencing, the degenerate MRE and a short flanking region were PCR amplified using the primers bi-dir-Miseq-F and bi-dir-Miseq-R. For cDNA and pDNA we used Phusion High-Fidelity PCR Master Mix with GC Buffer (NEB) and the following cycling conditions: initial denaturation (98°C for 30 s), 23 amplification cycles (98°C for 10 s, 65°C for 10 s, 72°C for 10 s) and final extension (72°C for 5 min). For cDNA from RNA recovered from polysome fractions we used KAPA HiFi HotStart ReadyMix (Fisher Scientific) and the following cycling conditions: initial denaturation (98°C for 30 s), 21 amplification cycles (98°C for 10 s, 65°C for 10 s, 72°C for 10 s) and final extension (72°C for 5 min). These initial PCR products were gel-purified using the QIAquick Gel Extraction Kit (Qiagen). We diluted the recovered product between 10 and 30 fold depending on band intensity.

To make amplicon libraries compatible with Illumina machines, we performed a second PCR to append TruSeq index sequences and p5/p7 adapters to each amplicon. We used a dual barcoding strategy where a unique combination of forward and reverse index primers were assigned to each biological sample. We performed the PCRs with Phusion High-Fidelity PCR Master Mix with GC Buffer (NEB) and the following cycling conditions: initial denaturation (98 °C for 30 s), 13 amplification cycles (98°C for 10 s, 62°C for 10 s, 72°C for 10 s) and final extension (72°C for 5 min). We used Agencourt AMPure XP beads (0.75X, Beckman Coulter) at 0.75X to purify the amplicon libraries which we subsequently quantified using the Qubit dsDNA HS Assay Kit (ThermoFisher Scientific). The samples were sequenced (150bp PE sequencing) on either the HiSeq4000 (Illumina) or the MiSeq v2 (Illumina).

### High-throughput sequencing data analysis

High-throughput sequencing data were analysed using R (Version 3.4.1) and all scripts are available upon request. After inspecting the quality of sequencing data with FastQC, we used the Biostrings package (version 2.44.2) to trim reads down to the MRE and subsequently count the occurrence of each type of variant of interest in all amplicon libraries. We calculated variant frequency by normalizing read counts of each variant of interest to total library read counts in the respective library. We calculated transcript abundance for each variant by dividing its read frequency in the cDNA library to its read frequency in the respective pDNA library. We calculated translation efficiency for variants present in polysome profiles by dividing their read frequency in the heavy-polysome-bound library by read frequency in the respective monosome-bound library.

### Validation of high-throughput sequencing results by RT-qPCR

To validate our high-throughput sequencing assay we randomly selected miR-17-MRE variants by screening colonies from a 1/30,000 dilution of the variant library by Sanger sequencing using primer bi-dir-MRE-seq-1. Colonies were screened until we identified 15 unique single and double nucleotide variants. These constructs were individually transfected into HEK-293-T cells in triplicate in addition to control reporters encoding a Cel-miR-67-MRE and a perfectly complementary miR-17 MRE using the PEI transfection method described above. 24 hours after transfection we extracted RNA using the RNeasy Mini Kit (Qiagen). For each replicate, cDNA was generated from 100 ng of total RNA using the QuantiTect Reverse Transcription kit (Qiagen). We performed RT-qPCR using the SsoAdvanced Universal SYBR Green Supermix kit (Bio-Rad) on a CFX384 real-time system (Bio-Rad) with primer pairs spanning the MRE (MRE_qPCR-F and MRE_qPCR-F) or within the iBlue transcript (iBlue_qPCR-F and iBlue_qPCR-F) which serves as an internal control. The ΔΔCt method was used to compare expression of all MRE variants to a Cel-67-MRE control reporter by comparing the Ct of ECFP to that of iBlue for each sample replicate.

### Lentiviral vector cloning and virus production

We generated PD-1 lentiviral expression vectors using standard restriction cloning methods. The parent vector AB.pCCL.sin.cPPT.GFP.miR-17-3p.sensor.PGK.dNGFR.WPRE was a gift from Brian Brown (Addgene plasmid #85866). To simplify subsequent cloning steps a SbfI recognition site was introduced downstream of the minimal CMV promotor. The human PD-1 ORF was amplified from the PD-1 BRET vector (a generous gift from Simon Davis) using the primers PD1_Lenti_Shuttle_F and PDL1_Lenti_Shuttle_R. We digested this PCR product, as well as the destination vector with SbfI and NheI and ligated them using T4 DNA ligase (NEB).

The In-Fusion HD Cloning System (Takara Clontech) was used to replace PD-1 in the lentiviral expression vector with cytoplasmic-localized ovalbumin (OVA) coupled to EGFP by a T2A peptide cleavage signal to create a OVA-T2A-EGFP vector. We PCR amplified T2A-EGFP from pX458 (A gift from Feng Zhang, Addgene plasmid #48138) with the primers GFP_in_fusion_F2 and EGFP-in-fusion-R. OVA (without the first 47 amino acids) was amplified from the OVACyt vector^34^ using the primers Ova_In_Fusion_R2 and Ova-in-fusion-F. We fused the parent vector (linearized with SbfI and NheI) with the two inserts following the In-Fusion manufacturer’s instructions. To create miSFIT-tuning vectors we generated MRE inserts from short oligonucleotides (IDT). MRE inserts were annealed and phosphorylated (T4 PNK, NEB) and introduced downstream of PD1 or OVA-T2A-EGFP by restriction cloning between NheI and AgeI. MRE insertion was confirmed by Sanger sequencing using primer BBBdir-seq-2.

To produce lentiviral particles in HEK-293T cells, we co-transfected each lentiviral transfer vector with pCMV-dR8.91 and pMD2.G at a ratio of 1.5:1:1 using Polyethylenimine (PEI, 1 mg/ml, Sigma-Aldrich) as described above. After 24 hours we exchanged the transfection media (DMEM, 2% FBS, no P/S) with full media (DMEM, 15% FBS). We collected and filtered (0.22 μm filter, Millipore) viral supernatant 24 hours later and stored it at −80°C until transduction. We transduced B16-F10 cells and Jurkat T-cells using un-concentrated viral supernatant. Jurkat T-cells (clone 1.G4) were maintained in RPMI-1640 media (Gibco) supplemented with 10mM HEPES (Life Technologies), 1mM Sodium Pyruvate (Life Technologies), and 15% Fetal Bovine Serum (FBS, GIBCO). B16-F10 melanoma cells were grown in DMEM (Gibco) supplemented with 15% FBS (Gibco).

### Flow cytometry and fluorescence-activated cell sorting

All flow-cytometry experiments were performed on the BD LSR Fortessa Analyzer or the FACSymphony (BD Biosciences) and data were analysed using FlowJo (Version 10.3.0). We harvested adherent cells (B16-F10 or HEK 293T) using 0.05% Trypsin with EDTA (Thermo Fisher Scientific). For experiments requiring antibody staining we washed cells with FACS buffer (PBS with 5% FBS) before and after staining. For an overview of our flow-cytometry gating strategies see Supplementary Fig. 3, 4, 5 and 6. To generate miSFIT cell lines we transduced cells at low multiplicity of infection (For Jurkat T-cells < 15% transduced, for B16-F10s < 3% transduced), waited 5 to 7 days and selected stably transduced cells by FACS using the SH800S cell sorter (SONY) with a 100μm sorting chip. We sorted pools of approximately 150,000 cells per line on the basis of NGFR expression.

### B16-F10 melanoma / T-cell co-cultures

To study how antigen levels influence T-cell activation and cellular fitness *in vitro* we co-cultured our B16-F10 OVA-miSFIT cell lines with OT-I T-cells. Primary splenocytes were harvested from C57BL/6, OT-I mice and stimulated with SIINFEKL peptide (20μg/mL Cambridge Peptides) and IL-2 (10 units/mL BioLegend) in RPMI-1640 (Gibco) supplemented with 10% FBS (Gibco), 1% P/S, (10,000 U/mL, Gibco), 10mM HEPES (Life Technologies), 1mM Sodium Pyruvate (Life Technologies), 50 μM 2-Mercaptoethanol (Gibco) and 1% MEM Non-Essential Amino Acids (Gibco). After 48 hours, CD8^+^ T-cells were isolated using the mouse CD8a^+^ T-cell Isolation Kit (Miltenyi Biotec). For melanoma fitness experiments, 20,000 B16-F10 cells (approx. 50/50 mixture of B16-F10 OVA-cells and OVA^+^, miSFIT cells) were seeded per well in a 96 well plate (n = 3 per cell line). After allowing B16-F10 cells to adhere for 3 hours, OT-I T-cells were added to each well at different T-cell: B16-F10 ratios. Mixed cultures were incubated overnight and analysed by flow cytometry. Relative fitness was calculated by dividing the frequency of NGFR^+^ cells in the +T-cell condition by the frequency of NGFR^+^ cells in the no-T-cell condition. For T-cell activation experiments, stimulated T-cells were rested for 72 hours prior to being co-cultured with individual OVA-miSFIT cell lines at an 8:1 T-cell to B16-F10 ratio (n = 5 per cell line). After 24 hours, we analysed T-cells by flow-cytometry. Antibody clones and suppliers are listed in Supplementary Table 2.

### *In vivo* tumour growth assays

For *in vivo* tumour growth assays, 150,000 B16-F10 OVA-miSFIT cells were intradermally injected into WT C57BL/6 recipient mice (n = 6 recipient mice per cell line) on day 0. We isolated OT-I T-cells and stimulated them for 48 hours (see above and Fig. 4a) and intravenously injected 500,000 CD8^+^, OT-I T-cells per recipient mouse on day 7. Following the T-cell infusion we measured tumours every second day using Digital Callipers (Fisher Scientific). The experimenter performing the measurement was blinded to tumour identity. We culled mice when tumours exceeded 95mm^2^ using approved methods. Mice that did not have detectable tumours by day 17 were excluded from the study.

For TIL analysis, tumours were injected as described above but OT-I T-cells were adoptively transferred on day 8 to reduce the likelihood of complete tumour clearance. On day 13 all mice were culled and spleens and tumours were harvested by dissection. Spleens were processed as described above and tumours were dissociated using the Tumor Dissociation Kit, mouse (Miltenyi Biotec). Cells were washed, blocked with TruStain fcX (Biolegend) and stained with antibodies as listed in Supplementary Table 2. Animal experiments were conducted under the constraints of a project licence approved by an internal Oxford review board and the UK home office.

## ACKNOWLEDGEMENTS

We thank Jamie Michaels and Noam Prywes for their comments on the manuscript. We thank Paul Sopp, Kevin Clark, Sally-Ann Clark and Craig Waugh from the WIMM Flow Cytometry Facility for providing training and technical support. We acknowledge Justin Deme and Kathryn Robson for their advice and technical assistance in performing polysome profiling experiments. M.K.T is supported by a Marie Skłodowska-Curie Individual Fellowship. Y.S.M is funded by the Clarendon Scholarship, the WIMM prize studentship and the Christopher Welch Fellowship. H.M.S is supported by The Royal Society of NZ, Catalyst Seeding grant. D.J.H.F.K. is funded by a CIHR Postdoctoral Fellowship. V.C. and M.B.B. are funded by the MRC, Cancer Research UK (CRUK Programme C399/A2291) and the Oxford National Institute for Health Research (NIHR). T.A.M is funded by Medical Research Council (MRC, UK) Molecular Haematology Unit Grant MC_UU_12009/6. T.A.B. was supported by a Radcliffe Department of Medicine/MRC Scholars Programme Studentship. T.A.F. was supported by MRC (G0902418), BBSRC (BB/N006550/1) and Wellcome Trust ISSF (105605/Z/14/Z).

## AUTHOR CONTRIBUTIONS

Y.S.M. conceived the study. Y.S.M., T.A.F and M.B.B designed the experiments and analysed the results. Y.S.M and M.B.B executed most of the experiments. H.B., T.A.B., M.K.T and V.A. helped with the experiments and provided technical expertise. U.G. performed I.V. injections. H.C-Y, M.F., H.M.S. and T.A.M. provided guidance and expertise. D.J.H.F.K provided guidance and assisted with data analysis. V.C. provided expertise and helped with the experimental design. Y.S.M. and T.A.F wrote the manuscript. All other authors provided feedback on the manuscript.

---

**Supplementary Figure 1.**
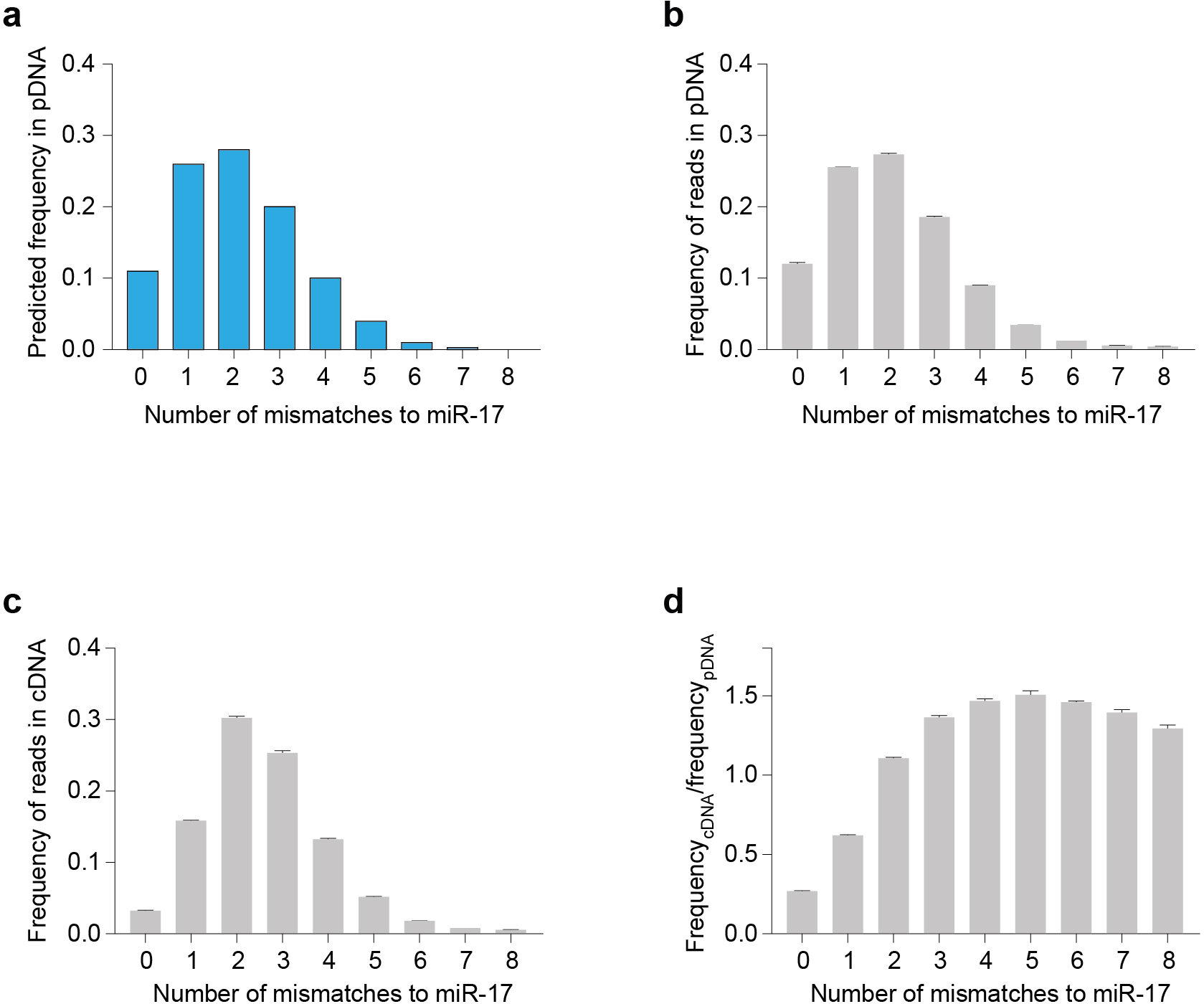
High-throughput study of the effect of mismatches on MRE functionality. **(a)** Expected probability distribution for the number of mismatches per MRE in the miR-17 MRE variant library (see Fig. 1a). **(b)** Frequency of reads containing between 1 and 8 mismatches to miR-17 in plasmid DNA (pDNA) isolated from HEK-293T cells transfected with the miR-17 MRE library (n = 3 biological replicates, mean +/− s.d.). **(c)** Frequency of reads containing 1 to 8 mismatches to miR-17 in cDNA libraries from the same pools of cells as in (b) (n = 3 biological replicates, mean+/− s.d.). **(d)** Average enrichment for MREs with varying numbers of mismatches calculated by dividing frequency of reads in cDNA libraries (c) by the frequency of reads in pDNA libraries (b).

**Supplementary Figure 2.**
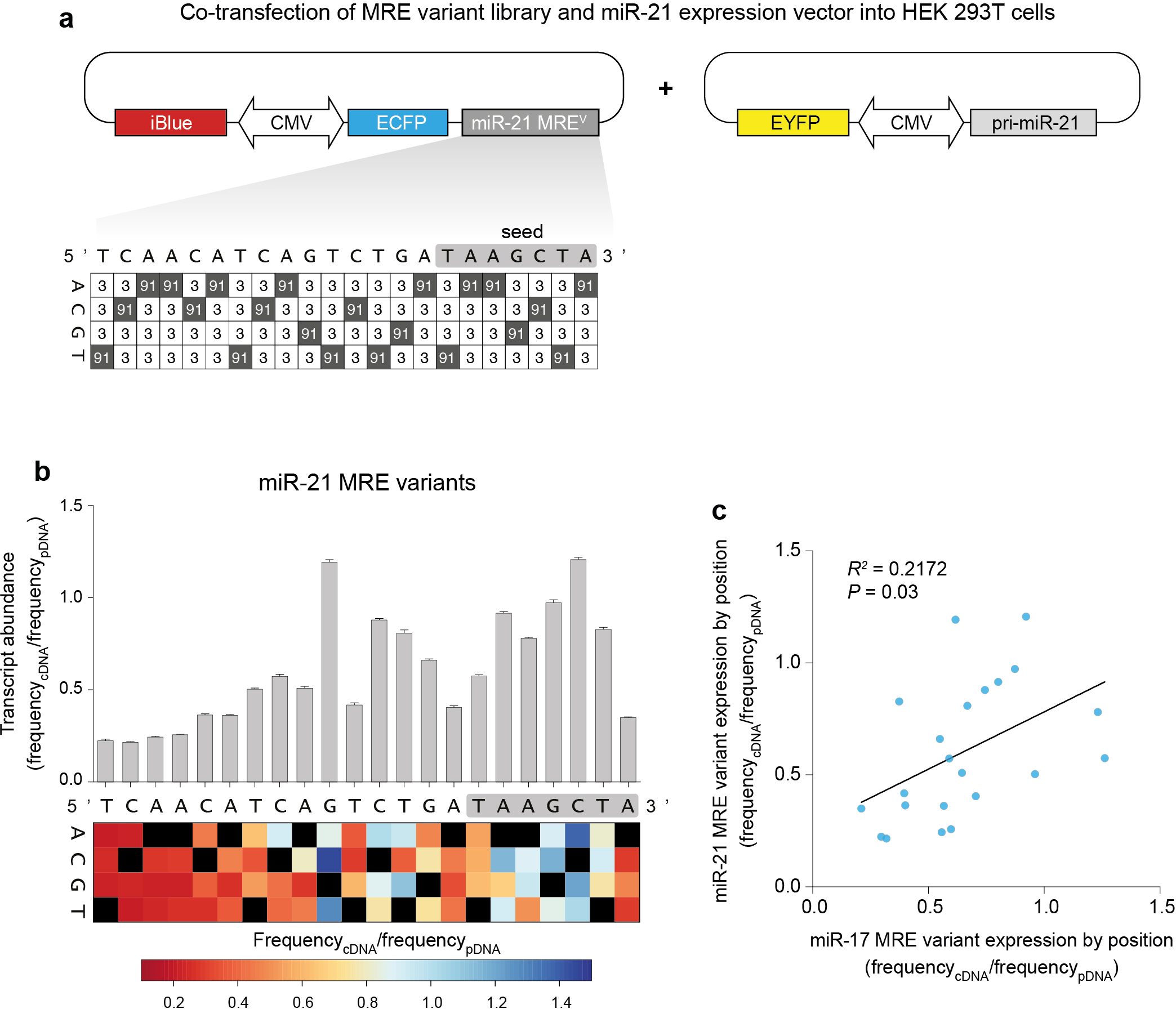
Single-nucleotide resolution analysis of the miR-21 MRE regulatory landscape. **(a)** Schematic representation of the miR-21 MRE library design (compare to miR-17 library in Fig. 1). Values indicate the proportion of nucleotides at each position in the MRE (shaded squares = nucleotides complementary to miR-21). Since miR-21 is expressed at low levels in HEK-293T cells, we overexpressed the primary miR-21 transcript using a plasmid vector which also delivers EYFP as a transfection control. **(b)** Impact of MRE variants on transcript abundance. Bar graph shows the impact of single nucleotide mismatches at each position in the MRE on the strength of reporter silencing by high-throughput sequencing (n = 3 biological replicates, mean+/− s.d.). Heat-map displays the effect of each possible mismatch by position and reflects the average of three replicates. (black squares = complementary bases) **(c)** Linear regression comparing the impact of mismatches at each position between the miR-17 and miR-21 synthetic MRE variants.

**Supplementary Figure 3.**
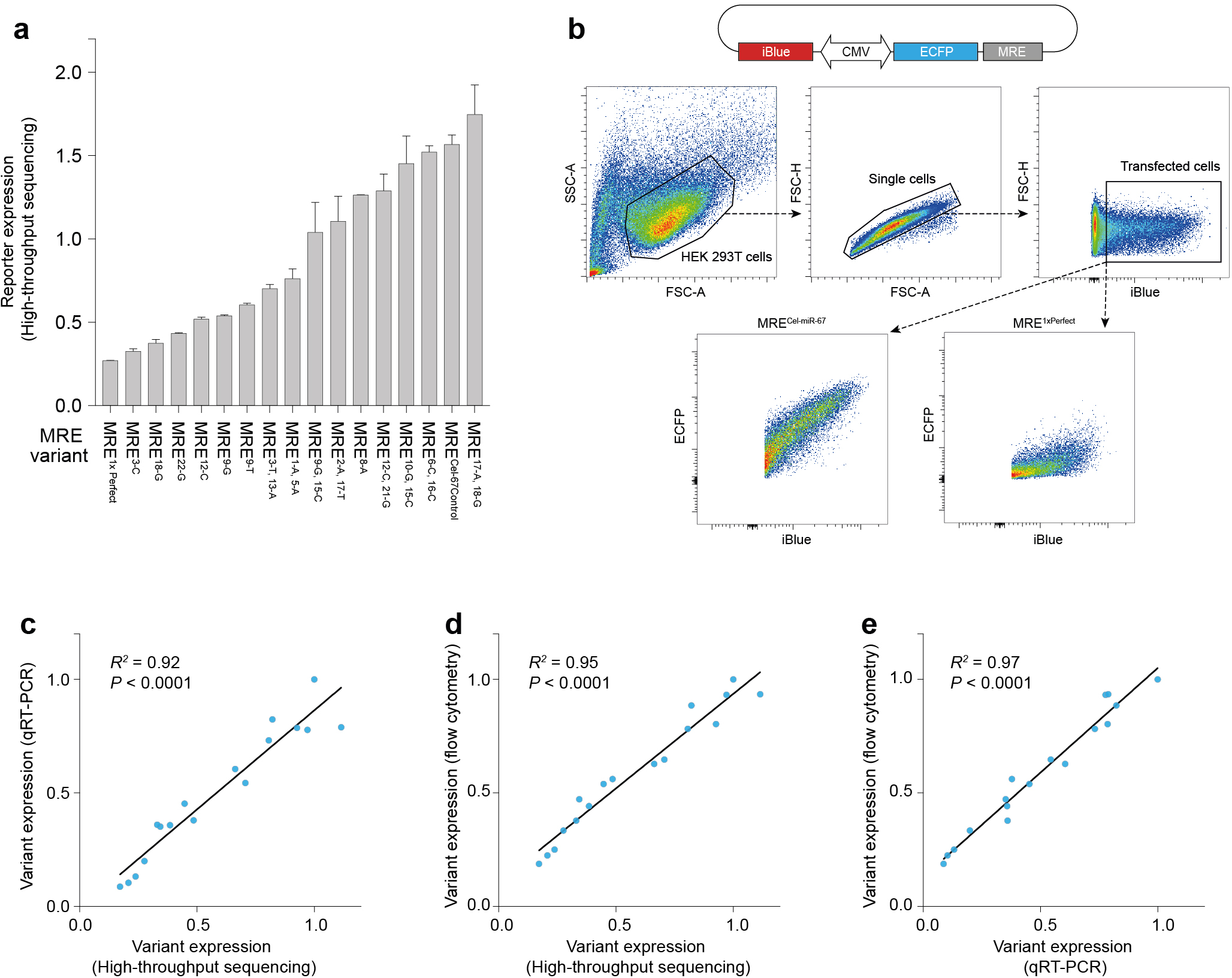
RT-qPCR and flow cytometry validate high-throughput MRE screen data. **(a)** Ranked impact of candidate MRE variants on transcript abundance. To select an unbiased set of miR-17 MREs from our variant library for validation using conventional methods, we transformed the plasmid library (see Fig. 1) into *E. coli* and randomly picked and screened colonies to identify 15 unique single or double variant MREs. Bar graph represents the effect on reporter expression of these variants relative to a perfect miR-17 MRE (MRE^1xPerfect^) and MRE^Cel-67 Control^, as determined by the high-throughput sequencing experiment (n = 3 biological replicates, mean+/− s.d.). **(b)** Flow cytometry validation gating strategy. To validate the accuracy of our high-throughput sequencing assay we independently transfected each of the 15 MRE variant reporters from the validation set and assayed ECFP expression by flow cytometry. **(c)** Linear regression comparing expression measured by high-throughput sequencing with expression measured by RT-qPCR in HEK-293T cells transfected with each of the 15 MRE variants in the validation set (*P* < 0.0001, slope differs from 0). We calculated expression by the LiiiCT method using iBlue as a reference gene. **(d)** Linear regression comparing variant expression measured by high-throughput sequencing and flow cytometry (flow cytometry expression was calculated by normalizing ECFP expression to iBlue on a single cell basis and taking the mean of that value for each MRE variant) (*P* < 0.0001, slope differs from 0). **(e)** Linear regression comparing expression as measured by RT-qPCR to flow cytometry (*P* < 0.0001, slope differs from 0).

**Supplementary Figure 4.**
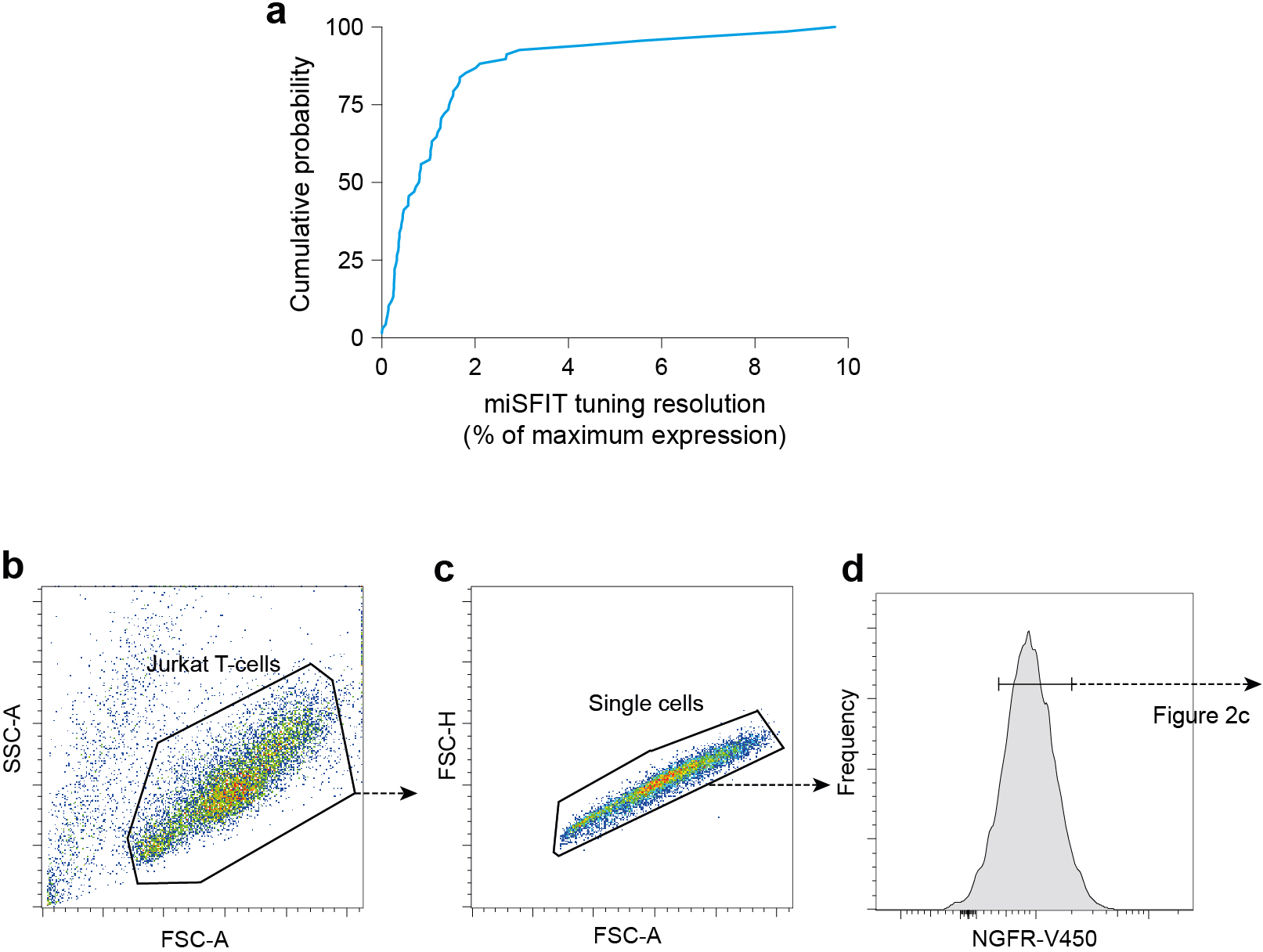
miSFITs enable precise tuning of gene expression. **(a)** Cumulative probability distribution of the precision that can be achieved using miSFIT technology. The distribution reflects the difference in ECFP expression between nearest single-nucleotide miR-17-MRE variants in HEK-293T cells. **(b - d)** Flow-cytometry gating strategy used to generate the data shown in Fig. 2c.

**Supplementary Figure 5.**
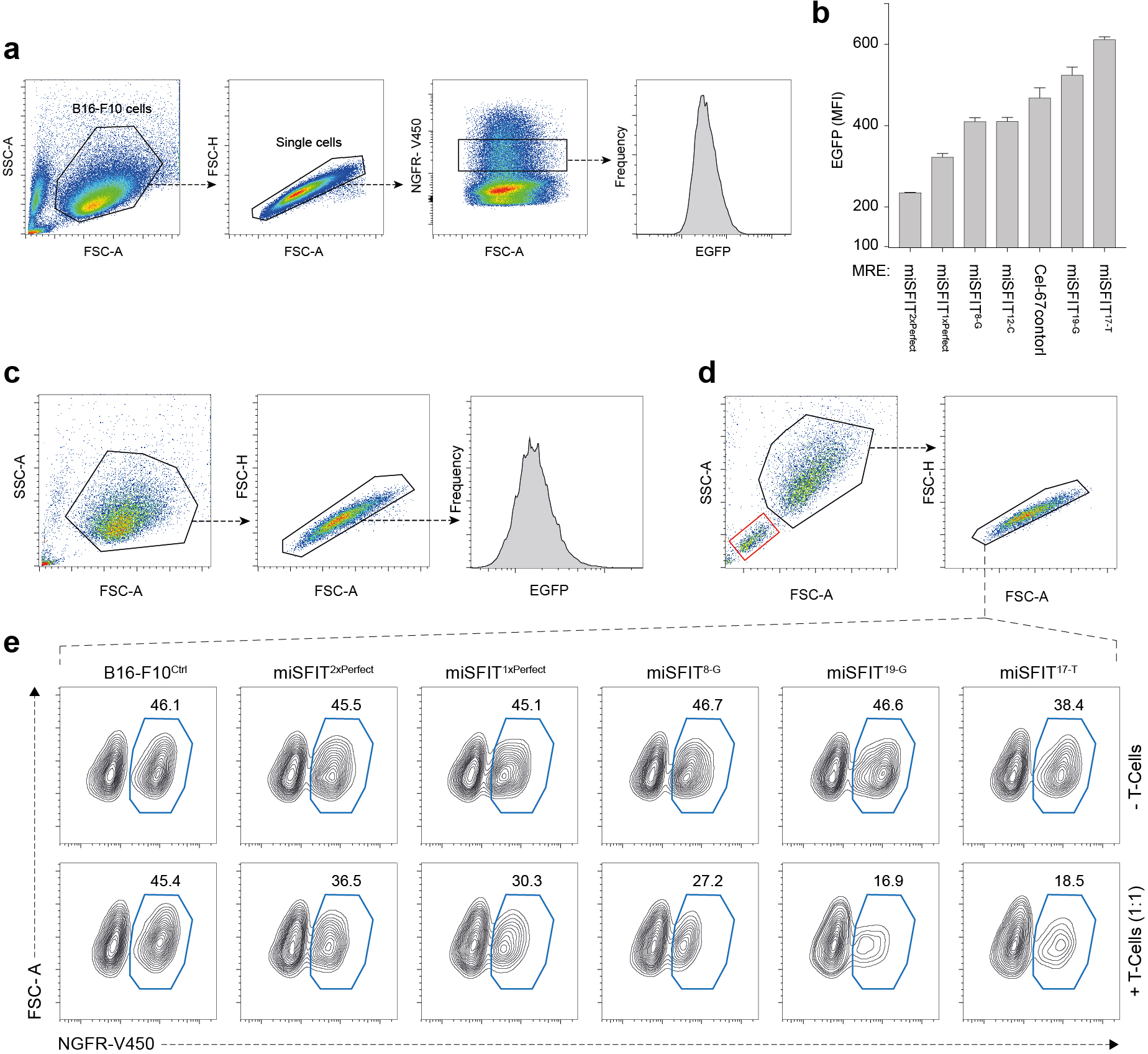
Flow cytometry analysis of OVA-T2A-GFP miSFIT cell lines. **(a)** Flow-cytometry gating strategy used to assess OVA-T2A-EGFP expression following transient transfection of seven different miSFIT variant constructs in B16-F10 cells. **(b)** EGFP expression of seven OVA-miSFIT constructs (n = 3 biological replicates, mean +/− s.d.). **(c)** Flow cytometry gating strategy used to analyse miSFIT variant cell lines following lentiviral transduction and cell sorting as shown in Figure 3b. **(d)** Flow cytometry gating strategy used for B16-F10 mixed co-culture experiments (red gate = T-cells). **(e)** NGFR expression following overnight mixed co-cultures of B16-F10 cell lines expressing each of five OVA-miSFIT variants as well as an OVA-negative control cell line with CD8^+^ OT-I T-cells at 1:1 T-cell: melanoma cell ratio (blue polygon gate = the percentage of NGFR^+^ (OVA-miSFIT) cells surviving after overnight selection; compare to a 3:1 ratio in Fig. 3e; the results of both experiments are summarized in Fig. 3f).

**Supplementary Figure 6.**
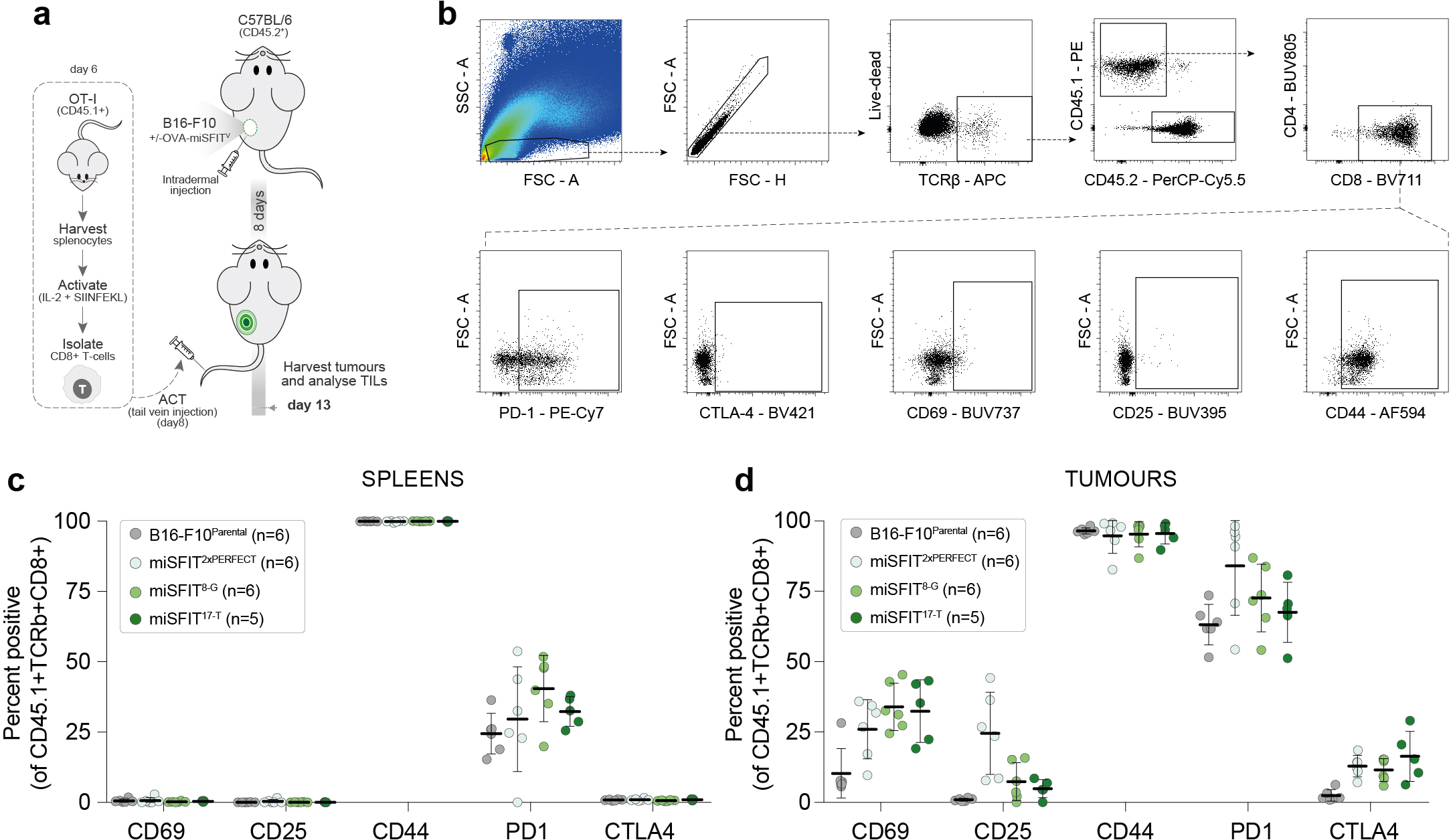
Analysis of Tumour Infiltrating Lymphocytes (TILs) **(a)** Schematic representation of the experimental design employed to assess the effect of OVA expression on Tumour Infiltrating Lymphocytes (Tlls). **(b)** Flow cytometry gating strategy used to analyse Tlls. Gates were drawn based on fluorescence-minus-one controls. OT-I donor mice are CD45.1^+^ while recipient mice are CD45.2^+^. **(c, d)** Cell surface marker expression on T-cells harvested from recipient mouse spleens **(c)** and tumours **(d)** (mean+/− s.d.).

## REFERENCES

1. Wei, F. et al. Strength of PD-1 signaling differentially affects T-cell effector functions. Proc Natl Acad Sci U S A 110, E2480–2489 (2013).

2. Eyquem, J. et al. Targeting a CAR to the TRAC locus with CRISPR/Cas9 enhances tumour rejection. Nature 543, 113–117 (2017).

3. Lee, T.I. & Young, R.A. Transcriptional regulation and its misregulation in disease. Cell 152, 1237–1251 (2013).

4. Alper, H., Fischer, C., Nevoigt, E. & Stephanopoulos, G. Tuning genetic control through promoter engineering. Proc Natl Acad Sci U S A 102, 12678–12683 (2005).

5. Mutalik, V.K. et al. Precise and reliable gene expression via standard transcription and translation initiation elements. Nat Methods 10, 354–360 (2013).

6. Salis, H.M., Mirsky, E.A. & Voigt, C.A. Automated design of synthetic ribosome binding sites to control protein expression. Nat Biotechnol 27, 946–950 (2009).

7. Keren, L. et al. Massively Parallel Interrogation of the Effects of Gene Expression Levels on Fitness. Cell 166, 1282–1294 e1218 (2016).

8. Ahler, E. et al. Doxycycline alters metabolism and proliferation of human cell lines. PLoS One 8, e64561 (2013).

9. Liberles, S.D, Diver, S.T, Austin, D.J. & Schreiber, S.L. Inducible gene expression and protein translocation using nontoxic ligands identified by a mammalian three-hybrid screen. Proc Natl Acad Sci U S A 94, 7825–7830 (1997).

10. Jackson, A.L. et al. Widespread siRNA “off-target” transcript silencing mediated by seed region sequence complementarity. RNA 12, 1179–1187 (2006).

11. Ferreira, J.P, Overton, K.W. & Wang, C.L. Tuning gene expression with synthetic upstream open reading frames. Proc Natl Acad Sci U S A 110, 11284–11289 (2013).

12. Wee, L.M, Flores-Jasso, C.F, Salomon, W.E. & Zamore, P.D. Argonaute divides its RNA guide into domains with distinct functions and RNA-binding properties. Cell 151, 1055–1067 (2012).

13. Mullokandov, G. et al. High-throughput assessment of microRNA activity and function using microRNA sensor and decoy libraries. Nat Methods 9, 840–846 (2012).

14. Vainberg Slutskin, I., Weingarten-Gabbay, S., Nir, R., Weinberger, A. & Segal, E. Unraveling the determinants of microRNA mediated regulation using a massively parallel reporter assay. Nat Commun 9, 529 (2018).

15. Guo, H., Ingolia, N.T, Weissman, J.S. & Bartel, D.P. Mammalian microRNAs predominantly act to decrease target mRNA levels. Nature 466, 835–840 (2010).

16. Meijer, H.A. et al. Translational repression and eIF4A2 activity are critical for microRNA-mediated gene regulation. Science 340, 82–85 (2013).

17. Eichhorn, S.W. et al. mRNA destabilization is the dominant effect of mammalian microRNAs by the time substantial repression ensues. Mol Cell 56, 104–115 (2014).

18. Landgraf, P. et al. A mammalian microRNA expression atlas based on small RNA library sequencing. Cell 129, 1401–1414 (2007).

19. Sharma, A. et al. Posttranscriptional regulation of interleukin-10 expression by hsa-miR-106a. Proc Natl Acad Sci U S A 106, 5761–5766 (2009).

20. Peterson, S.M. et al. Common features of microRNA target prediction tools. Front Genet 5, 23 (2014).

21. Iwakawa, H.O. & Tomari, Y. The Functions of MicroRNAs: mRNA Decay and Translational Repression. Trends Cell Biol 25, 651–665 (2015).

22. Floor, S.N. & Doudna, J.A. Tunable protein synthesis by transcript isoforms in human cells. Elife 5 (2016).

23. Baumeister, S.H, Freeman, G.J, Dranoff, G. & Sharpe, A.H. Coinhibitory Pathways in Immunotherapy for Cancer. Annu Rev Immunol 34, 539–573 (2016).

24. Farkona, S., Diamandis, E.P. & Blasutig, I.M. Cancer immunotherapy: the beginning of the end of cancer? BMC Med 14, 73 (2016).

25. Buus, S. et al. Sensitive quantitative predictions of peptide-MHC binding by a ‘Query by Committee’ artificial neural network approach. Tissue Antigens 62, 378–384 (2003).

26. Lever, M. et al. Architecture of a minimal signaling pathway explains the T-cell response to a 1 million-fold variation in antigen affinity and dose. Proc Natl Acad Sci U S A 113, E6630–E6638 (2016).

27. Huang, J. et al. A single peptide-major histocompatibility complex ligand triggers digital cytokine secretion in CD4(+) T cells. Immunity 39, 846–857 (2013).

28. Schroder, A.R. et al. HIV-1 integration in the human genome favors active genes and local hotspots. Cell 110, 521–529 (2002).

29. Hogquist, K.A. et al. T cell receptor antagonist peptides induce positive selection. Cell 76, 17–27 (1994).

30. Schreiber, R.D, Old, L.J. & Smyth, M.J. Cancer immunoediting: integrating immunity’s roles in cancer suppression and promotion. Science 331, 1565–1570 (2011).

31. Zehn, D., Lee, S.Y. & Bevan, M.J. Complete but curtailed T-cell response to very low-affinity antigen. Nature 458, 211–214 (2009).

32. Brown, B.D. et al. Endogenous microRNA can be broadly exploited to regulate transgene expression according to tissue, lineage and differentiation state. Nat Biotechnol 25, 1457–1467 (2007).

33. Michot, J.M. et al. Immune-related adverse events with immune checkpoint blockade: a comprehensive review. Eur J Cancer 54, 139–148 (2016).

34. Rowe, H.M. et al. Immunization with a lentiviral vector stimulates both CD4 and CD8 T cell responses to an ovalbumin transgene. Mol Ther 13, 310–319 (2006).

